# An in-silico study on SARS-CoV-2: Its compatibility with human tRNA pool, and the polymorphism arising in a single lineage over a month

**DOI:** 10.1101/2020.07.23.217083

**Authors:** Manish Prakash Victor, Rohit Das, Tapash Chandra Ghosh

**Author notes:** To whom correspondence should be addressed., Correspondence may also be addressed to Dr. Tapash Chandra Ghosh.

## Abstract

SARS-CoV-2 has caused a global pandemic that has costed enormous human lives in the recent past. The present study is an investigation of the viral codon adaptation, ORFs’ stability and tRNA co-adaptation with humans. We observed that for the codon usage bias in viral ssRNA, ORFs have near values of folding free energies and codon adaptation index with mRNAs of the human housekeeping CDS. However, the correlation between the stability of the ORFs in ssRNA and CAI is stronger than the mRNA stability and CAI of HKG, suggesting a greater expression capacity of SARS-CoV-2. Mutational analysis reflects polymorphism in the virus for ORF1ab, surface glycoprotein and nucleocapsid phosphoprotein ORFs. Non-synonymous mutations have shown non-polar substitutions. Out of the twelve mutations nine are for a higher t-RNA copy number. Viruses in general have high mutation rates. To understand the chances of survival for the mutated SARS-CoV-2 we did simulation for synonymous mutations. It resulted in 50% ORFs with higher stability than their native equivalents. Thus, considering only the synonymous mutations the virus can exhibit a lot of polymorphism. Collectively our data provides new insights for SARS-CoV-2 mutations and the human t-RNA compatibility.

**Significance:** Survivability of SARS-CoV-2 in humans is essential for its spread. It has overlapping genes exhibiting a high codon optimization with humans even after a higher codon usage bias. They seem to possess cognizance for high copy number t-RNA (cognate or near-cognate) in humans, while mutating. Even though, it has been well established that native transcripts posses the highest stability, our in-silico studies show that SARS-CoV-2 under mutations give rise to ORFs with higher stability. These results significantly present the virus’s ability and the credibility of survival for the mutants. Despite its focus on a geographical location it explains the ongoing behavior of SARS-CoV-2 for a steady existence in humans as all the different lineages have a common origin. Wuhan, China.

## 1. Introduction

According to the world health organization the first human case of covid-19 was reported on 31^st^ December, 2019 in the Wuhan province of China. Since then, within a span of seven months the virus has spread across the globe infecting ~ >14 million and killing over ~608,000 people. Ministry of health and family welfare (Govt. of India) reported ~>402,529 active cases and ~>28,084 deaths as on 21^st^ of july, 2020. COVID-19 causes severe acute respiratory illness and in some cases acute respiratory distress syndrome (*1*). Additionally, COVID-19 is seen to cause systemic inflammation leading to sepsis, heart failure and multi-organ dysfunction(*2*). Covid-19 is a viral disease caused due to the virus SARS-CoV-2 which has a close similarity with SARS-CoV and MERS-CoV(*3, 4*).

The genome organization of SARS-CoV-2 shows a single-stranded positive sense RNA (*5*). It has twelve functional open reading frames containing ~30,000 nucleotides which encodes for ~7096 residues long polyprotein (*6*). They represent all the structural and the non-structural proteins that help the virus to infect and survive in the host’s cell (*7*). The open reading frames are ORF1ab consisting of overlapping ORF1a and ORF1b, followed by other proteins i.e. Spike (S), ORF3a, Envelope (E), Membrane (M), ORF6, ORF7a, ORF7b, ORF8, Nucleocapsid (N), and ORF10(*8*). After a virus has infected the host, it overtakes the host’s translational machinery by attaching the RNA to the ribosomes which turns the host’s cells into a ribovirocell. In a ribovirocell the viral and the cellular genome co-exist and the cell still divides producing virions (*9, 10*).

Since, viruses depend on their host for its gene translation; our study provides a comparative analysis on the codon usage overlap of the viral genome with the human housekeeping genes. We have also tried to study its compatibility with the human cognate and near-cognate t-RNA pool. While housekeeping genes have a uniform and high expression in all the cells of an organism they provide a better insight for a comparative analysis on the codon usage.

A codon is the combination of three out of four nucleotides Adenine, Thymine (Uracil in RNA), Cytosine and Guanine. There are a total of 61 codons that encodes 20 amino-acids. Leaving aside the amino-acids tryptophan and methionine all the other 18 amino-acids are encoded by two to six degenerate codons. The degenerate codons are usually referred as the synonymous codons due to their property of encoding the same amino-acid. Along-with encoding the amino-acids, codons are helpful in carrying extra information (*11*). Like the specific arrangements of nucleotides have been seen to act as recognition sites for protein interaction with DNA/RNA, modulate mRNA stability controlling their secondary structures which in-turn guides the ribosomal abundance during translation (*12, 13*). Hence, genes/genomes in different organisms tend to show the usage of a subset of codons from the pool of 61 codons depending upon their intrinsic requirements. Such a phenomenon is known as codon usage bias. Generally, higher codon usage bias is seen in highly expressed genes (*14, 15*). Also, these set of genes are strongly optimized to a small set of t-RNAs (*16*). According to the translational selection hypothesis the aforementioned conditions are suitable for rapid translation with higher efficiency and fidelity (*17, 18*). We found that the previous and all the mutated SARS-CoV-2 has comparatively higher codon usage bias than the human housekeeping genes and is proficiently compatible with the human t-RNA pool of anti-codons.

In general, viruses have mutation rates which is almost a million times higher than their hosts (*19*). So to understand the mutational capacity aiding in survival for SARS-CoV-2 we performed multiple simulations of synonymous mutations on each ORF. The result reveals that under the inherent codon usage bias the virus can attain ORF stability maxima greater than their native sequences which will be helpful in greater translational efficiency(*13*). Hence, SARS-CoV-2 still poses a threat due to its immense mutational impetus that has the ability to keep the bias and anti-codon compatibility intact, and produce stable ORFs that can result in translational efficiency to be greater than its host.

## 2. Materials and methods

### 2.1. Data collection

We downloaded all the CDS for SARS-CoV-2 and *Homo sapiens* housekeeping genes from the NCBI database (Refer Table S2.1, Supplementary tables). Longest transcripts were selected. MT539164 is a SARS-CoV-2 strain is referred as the predecessor strain in our study; sequences collected on a later date are referred as successor strains. Copy number for the *Homo sapiens* tRNA molecules containing anticodons for each 61 codons were obtained from GtRNAdb 2.0 release 18.1(*20*). The near-cognate t-RNA mappings were done for the unavailable t-RNA containing anti-codons with respect to codons in SARS-CoV-2 (Refer Table S2.2, Supplementary tables) using the dataset available from (*21*).

### 2.2. Randomization of the SARS-CoV-2 ORFs

Mutations were simulated through synonymous codons’ randomization within the gene without affecting the intrinsic codon usage frequency and amino-acids. (Refer S1.1, supplementary materials and methods).

### 2.3. Codon adaptation index and co-adaptation index calculations (co-AI)

Emboss cusp, cai and chips programs were used to find the codon usage frequency, codon adaptation index and effective number of codons (*22*). co-AI was calculated as done by Victor et al. (*23, 24*).

### 2.4. Folding energy calculation

The folding free energies of ORFs were calculated for windows of size 50 nucleotides and a step size of 30 nucleotides using seqfold (*25*). Energies predicted are averaged over the number of segments to obtain the folding free energy for an entire ORF (Refer S1.2, supplementary materials and methods).

### 2.4 þn, þs and Statistical calculations

We have used Pearson correlations with 95% level of confidence as a measure of significance unless otherwise stated. R language and environment (https://www.r-project.org/) was used to perform statistical analyses. For understanding the choice of statistical method on small samples (Refer S1.3, supplementary material and methods). Python package dnds_cal-0.0.2 with attribute for polymorphism (pn, ps) calculations was used.

## 3. Results

### 3.1 ENC, co-AI and CAI of SARS-CoV-2 with human (HKG)

Effective number of codons (ENC) quantifies the codon usage bias in genes/genomes. It ranges from 20(extremely biased) to 61(nil bias). Thus, a higher value of ENC indicates lower bias and vice-versa. In our dataset for SARS-CoV-2 predecessor strain the ENC=45.001 and for Housekeeping genes the ENC=56.550. Other successor strains have also shown very near values of ENC amongst themselves and with the predecessor (Table1). co-adaptation index (co-AI) or tRNA co-adaptation is an indicator of the codon optimization to the tRNA pool (*23, 24*). As viruses depend on the translational machinery of the host, their inherent codon composition needs to be compatible with the host’s tRNA pool. The co-AI for SARS-Cov-2 (Pearson ρ=0.304; P<0.05; n=61; <u>MT539164</u>) and HKG (Pearson ρ=0.350; P<0.05; n=61) shows a significant positive correlation. Successor strains have shown very near values of co-AI amongst themselves and with the predecessor strain (Table1). Codon adaptation index (CAI) measures the relative adaptiveness of the codon usage of a gene towards the codon usage of highly expressed genes, and it is used to estimate the gene expression level (*24*). Using the codon set of highly expressed genes in *Homo sapiens*, SARS-CoV-2 predecessor strain yielded CAI _mean_ = 0.637 which is very close to HKG CAI _mean_ = 0.73 (Refer Table S2.3, Supplementary tables). Successor strains have shown identical CAI values amongst themselves and with the predecessor strains.

**Table1:**
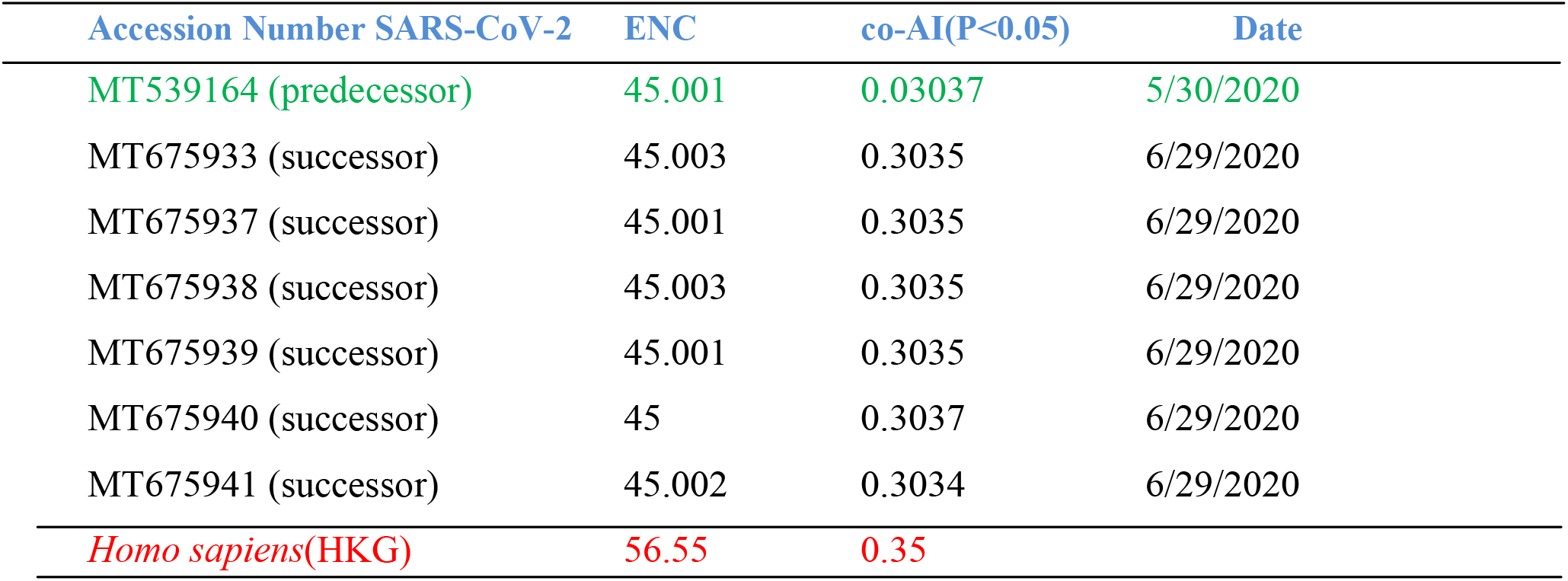
Effective number of codons (ENC) and co-adaptation index (co-AI) for different viral strains and *Homo sapiens* (HKG). HKG show a lower codon usage bias but higher codon optimization

### 3.2 ORF stability and gene expression level in SARS-CoV-2

Here, we have considered the structural stability of the ORFs, owing to its secondary structure. ORF stability referred as the minimum folding energy (mFE) is measured by Gibbs free energy; with greater negative values indicating higher feasibility in folding, thus, higher ORF stability (Refer Table S2.3, Supplementary tables). There is a significant negative correlation between CAI and mFE for the predecessor strain (Pearson ρ = −0.63; P <0.05; n=12). A similar trend is seen for the house keeping genes (Pearson ρ = −0.563; P <0.1; n=12) (Refer S1.3, Supplementary materials and methods on the selection of statistical methods for small samples). It indicates that stable ORFs have higher expression values and SARS-CoV-2 exhibits a stronger correlate demonstrating the virus’s innate ability to have a greater expression capacity than humans.

### 3.3 Simulating mutations: synonymous random shuffling within SARS-CoV-2 ORFs

Hundred in-silico mutated sequences were generated through random synonymous codon shuffling of each ORF in the predecessor strain. For each random sequence of a gene we maintained the inherent codon usage frequency and the amino-acid sequences equivalent to each gene in the predecessor strain. Minimum folding energy for all the sequences were calculated (Refer S1.1, Supplementary materials and methods). We found that 50% of the ORFs showed stability maxima, higher than their native ORFs (Refer Table S2.4, Supplementary tables). The difference between the stabilities obtained is significant which was found through the one-tailed t-test (t= −1.618; P<0.1; n=12)

### 3.4 Evaluation of SARS-CoV-2 mutations

Analyzing the six different strains (successor strains) of the SARS-Cov-2 (dated: 2020-05-30 & 2020-06-29) we have observed changes in ORF1a polyprotein, ORF1b polyprotein, surface glycoprotein and nucleocapsid phosphoprotein. They have incorporated synonymous as well as non-synonymous mutations (Refer Table S2.5, Supplementary tables). Overall twelve codon changes (different frequencies) have been recorded, resulting in a miniscule GC-content reduction (~0.01%) in entirety (Table 2). All the non-synonymous changes are non-polar (Table 3). Out of the twelve mutations recorded, seven showed the choice for rare codons, which after near-cognate t-RNA mapping has shown the choice for higher t-RNA gene copy number, three showed the choice for frequently used codons and the rest two are neutral changes. In totality nine mutations are for the higher gene copy number of cognate and near-cognate t-RNA containing the anticodons (Table 2). The polymorphism was calculated between the predecessor strain and all the successor strains as the ratio of non-synonymous to synonymous polymorphisms (þn/ þs). þn/ þs for all the mutations has generated values >1 (Table 4).

**Table 2:** t-RNA copy number change due to the mutations in SARS-CoV-2 and the near-cognate t-RNA mapping for the changed codons. zero shows the absence of cognate t-RNA

| Codon Change | t-RNA copy number change | t-RNA mapping after change |
| --- | --- | --- |
| CAG ⇒ CCG | 13 ⇒ 04 | |
| TGC ⇒ TGT | 29 ⇒ 00 | 29 ⇒ 29 |
| GTG ⇒ GTT | 11 ⇒ 09 | |
| TTT ⇒ TTC | 00 ⇒ 10 | |
| CTT ⇒ CCT | 09 ⇒ 09 | |
| CTT ⇒ TTT | 09 ⇒ 00 | 09 ⇒ 10 |
| TTA ⇒ TTG | 04 ⇒ 06 | |
| CTC ⇒ TTC | 00 ⇒ 10 | |
| TTG ⇒ TTT | 06 ⇒ 00 | 06 ⇒ 10 |
| GAC ⇒ GAT | 13 ⇒ 00 | 13 ⇒ 13 |
| TCA ⇒ TTA | 04 ⇒ 04 | |
| ACA ⇒ ATA | 06 ⇒ 05 | |

**Table 3:**
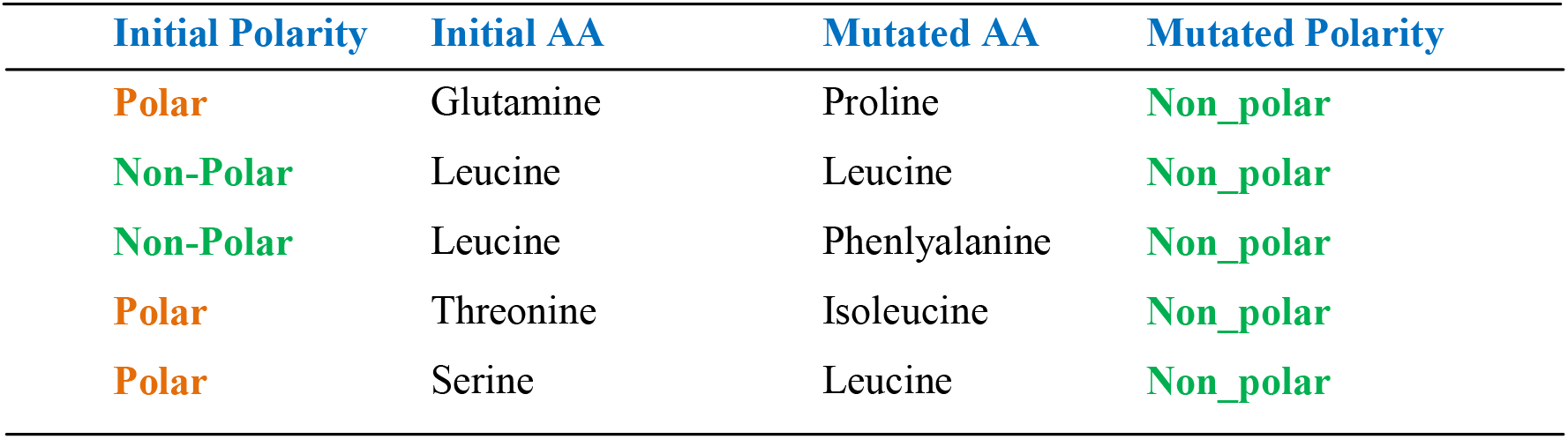
Non-synonymous mutations in SARS-Cov-2 have shown a preference for Non-polar amino-acids (AA)

| Initial Polarity | Initial AA | Mutated AA | Mutated Polarity |
| --- | --- | --- | --- |
| **Polar** | Glutamine | Proline | **Non_polar** |
| **Non-Polar** | Leucine | Leucine | **Non_polar** |
| **Non-Polar** | Leucine | Phenlyalanine | **Non_polar** |
| **Polar** | Threonine | Isoleucine | **Non_polar** |
| **Polar** | Serine | Leucine | **Non_polar** |

**Table 4:**
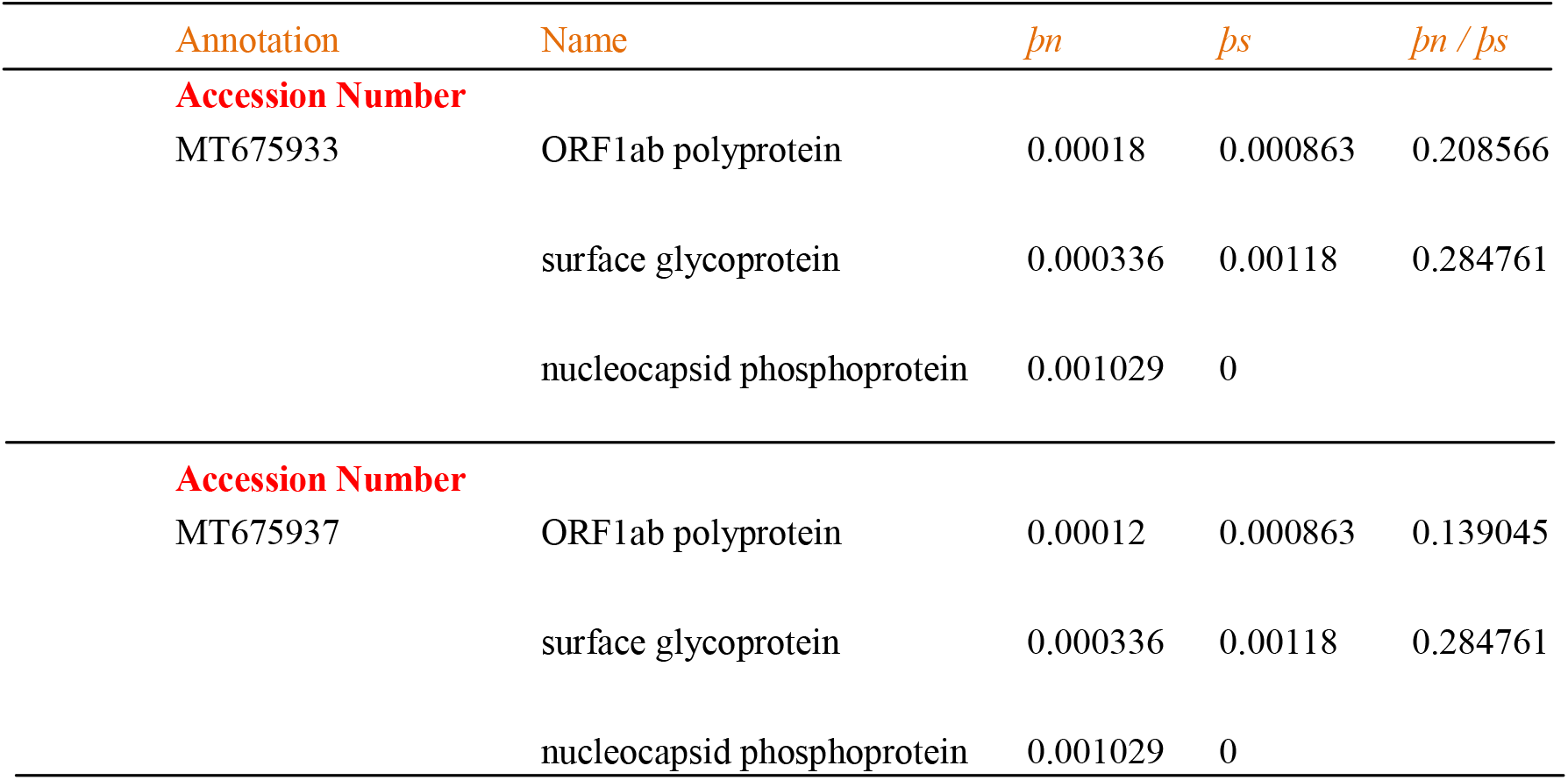
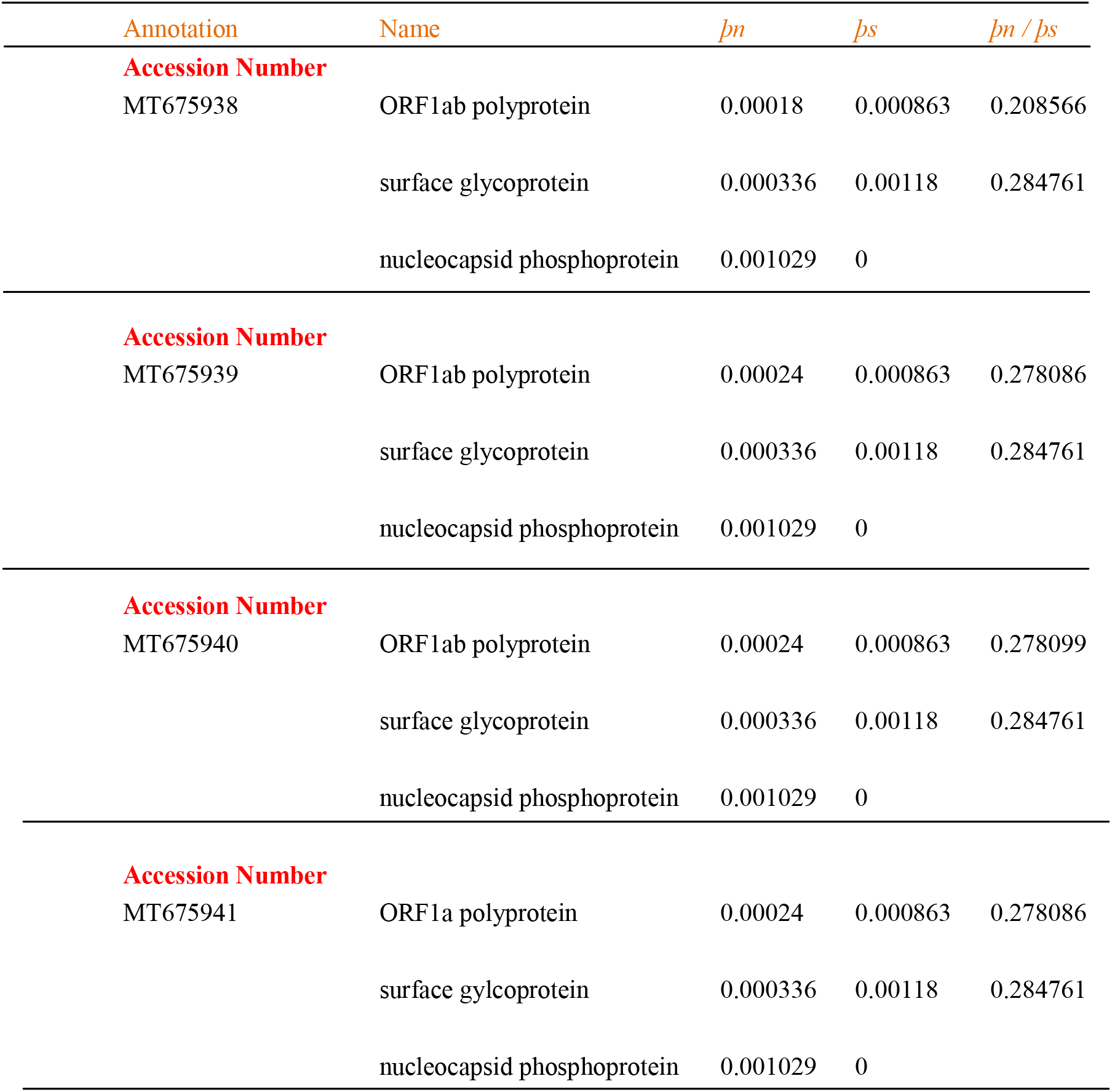
Synonymous and non-synonymous polymorphisms in SARS-CoV-2. To investigate the selection within a species we calculate it by *þn / þs* because the nucleotide changes are polymorphism which is segregating within a population. They are not fixed differences between lineages

## 4. Discussions

### 4.1 Analyzing the mutations in SARS-CoV-2

RNA viruses have an extremely high mutation rates. It can be around a million times higher than their host. These mutations account for their heightened virulence and evolvability. Higher mutation rates are a consequence of selection of variants exhibiting faster genomic replication (*19*). In a month’s time, SARS-CoV-2 has shown mutations in the ORF1ab polyprotein, surface glycoprotein and nucleocapsid phosphoprotein. The former two proteins have been shown to play an important role in the pathogenecity of the virus and the latter is responsible for the ssRNA packaging (*6, 26*). All of the mutations which are non-synonymous incorporated non-polar amino-acids in their protein (Table 3). Earlier analysis on the amino-acid composition of proteins have shown that non-polar amino-acids facilitate in the initiation and the propagation of protein folding (*27*), suggesting that the mutations here, might be required for improved protein folding that may provide enhanced proliferation for the virus particles. The mutations occurring in successor strains of a species (generally for viruses) are quantified as the ratio of the polymorphisms occurring in them. þn and þs calculates the relative abundance of non-synonymous and synonymous polymorphism. Diversifying or positive selection happens when þn/þs>1, whereas þn/þs<1 suggests purifying selection (*28, 29*). Looking at the values of þn/þs (Table 4, Figure 1) obtained in our dataset it is indicative of a purifying selection for the mutations. Even though genes have shown a purifying selection in a month’s time, the possibilities of further mutations cannot be discarded due to the inherently higher mutation rates of the viruses in general(*19*) and the value is not suggestive of the fixation of the present strains(*30*).

**Figure 1:**
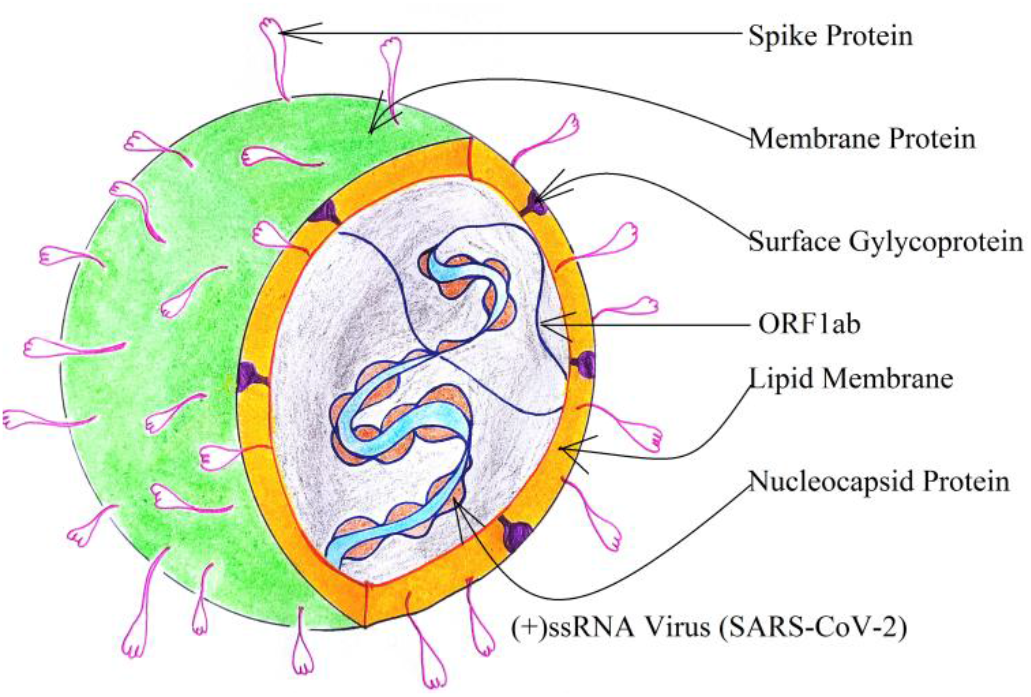
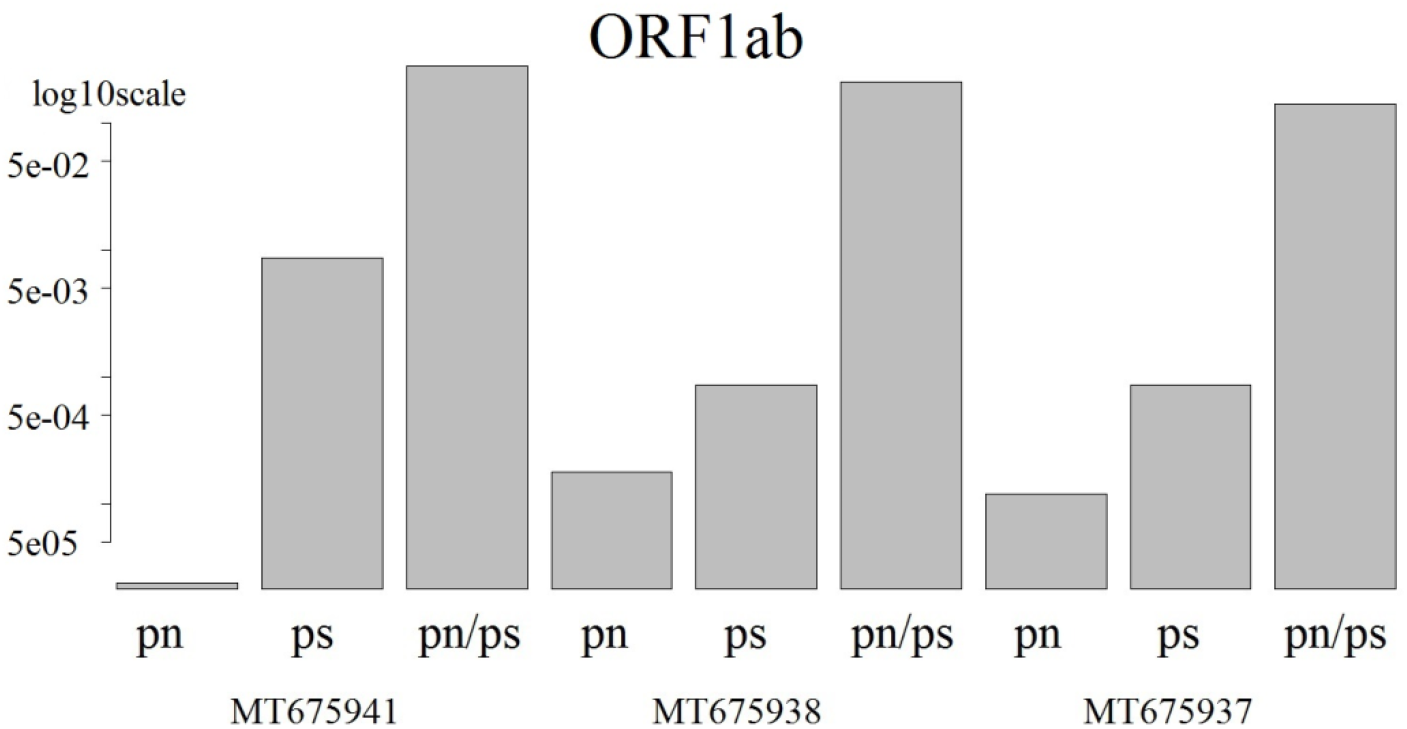
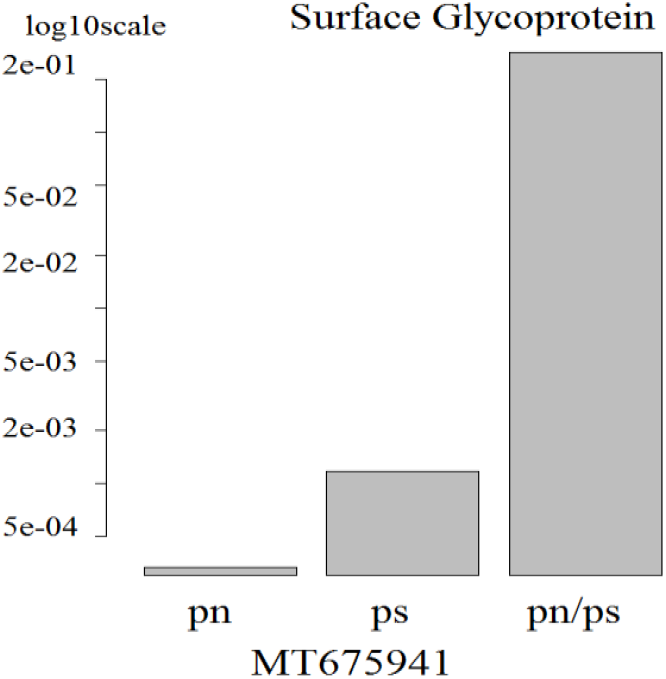
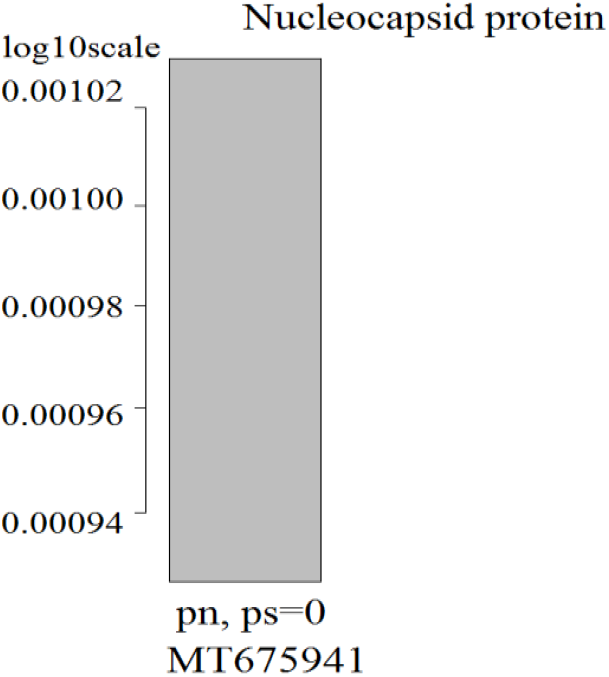
Schematic representation of the virus showing various polymorphisms in different strains. Overlapping values for the different strains have been omitted Figure 1.1: Schematic representation of the virus SARS-CoV-2 Figure 1.2: Polymorphism in the ORF1ab. Figure 1.4: Polymorphism in the nucleocapsid protein

### 4.2 Homo sapiens’ housekeeping genes and its importance in SARS-CoV-2 gene evaluation

Housekeeping genes are established to be evolutionarily conserved set of genes that tends to have higher codon adaptation index. They are required for the basic cell maintenance processes. Hence, they exhibit uniform expression levels throughout different cells under different conditions (*31*). For a comparative analysis in our further studies on simulating mutations, t-RNA compatibility and gene expression with SARS-CoV-2 we curated a set of human housekeeping genes (Refer Table S2.1, Supplementary tables). As SARS-CoV-2 affects the respiratory system we needed a set of housekeeping genes that can be relied upon even if there was any kind of pulmonary infection. The set of housekeeping genes (HKG) were taken as a control as suggested through the acute pulmonary inflammation study by(*32*).

### 4.3 Anti-codon mapping in SARS-CoV-2

Albeit humans have nearly five hundred different t-RNA genes but only forty eight anticodons are represented by them (*33*). However, humans use all the sixty one codons in their genome. This clearly shows two important aspects in the translational machinery: (i) for some codons, translation is achieved through near-cognate t-RNAs; (ii) there are multiple copies of the same t-RNA. Due to the dependence of the virus on its host for translation, the absentee cognate t-RNAs will be replaced by a near-cognate t-RNA (*21*). Hence, for a proper calculation of the co-adaptation index the codons lacking cognate t-RNA were needed to be mapped with the most appropriate near-cognate t-RNA in humans (Refer Table S2.2, Supplementary tables). The mapping is suggestive of the availability of the cognate and near-cognate t-RNAs for all the codons during translation(*21*). Moreover, it creates a clearer picture about the choices of near-cognate t-RNAs by SARS-CoV-2, and in our dataset we found that the choices are towards either highest or moderately higher copy number of near-cognate t-RNA containing anti-codons.

### 4.4 Codon usage bias, t-RNA adaptation, and gene expression of SARS-CoV-2

Effective number of codons is the quantification of the codon usage bias for a gene or a genome indicating the subset of codon usage from the pool of available 61 codons(*34*). But, it doesn’t tell about either the nucleotide composition or codon usage overlap in a comparative genome analysis. The ENC for HKG is higher to SARS-CoV-2 (Refer Results). Thus, SARS-CoV-2 has a more biased gene compared to the housekeeping genes. Generally, a biased gene tends to show a higher gene expression level compared a lowly biased system (*18, 35, 36*). In order to check the same, and codon usage overlap we calculated the CAI (a proxy for gene expression) for SARS-CoV-2 (Refer Results). As the virus relies on the host’s molecular machinery we assumed the human codon reference set of highly expressed genes for the calculation of CAI (Refer Results). Thus, CAI is helpful in understanding the overlap of codon usage patterns. The SARS-CoV-2 CAI value showed an 85.36% overlap in the expression magnitude with the housekeeping genes (Figure 2). In fact a biased system cannot always show a high gene expression level unless codon optimization is present(*37*). For a biased gene this values is an indicator of a sufficient codon optimization with the host. Even though CAI values are indicative of the gene expression levels it’s intuitive to think on the role of t-RNA abundance in influencing the gene expression (i.e. protein abundance). We calculated the co-AI, a correlate between the codons and t-RNA gene copy number (Refer Results). co-AI for HKG is 0.350 and predecessor strain is 0.304 (Refer Results, Table 1). The co-AI values were also calculated for the successors. It was found that all the successors have approximately the same values and is very near to the predecessor strain (Table 1). This strongly confirms a very close proximity of the codons in SARS-CoV-2 with the human t-RNA pool. The results signify that the codon composition of the SARS-CoV-2 is highly optimized with the anticodons in *Homo sapiens.* Here, it can be propounded that even under the mutations SARS-CoV-2 shows high codon optimization, hence, during its infection in the host the expression might be higher than other host genes.

**Figure2:**
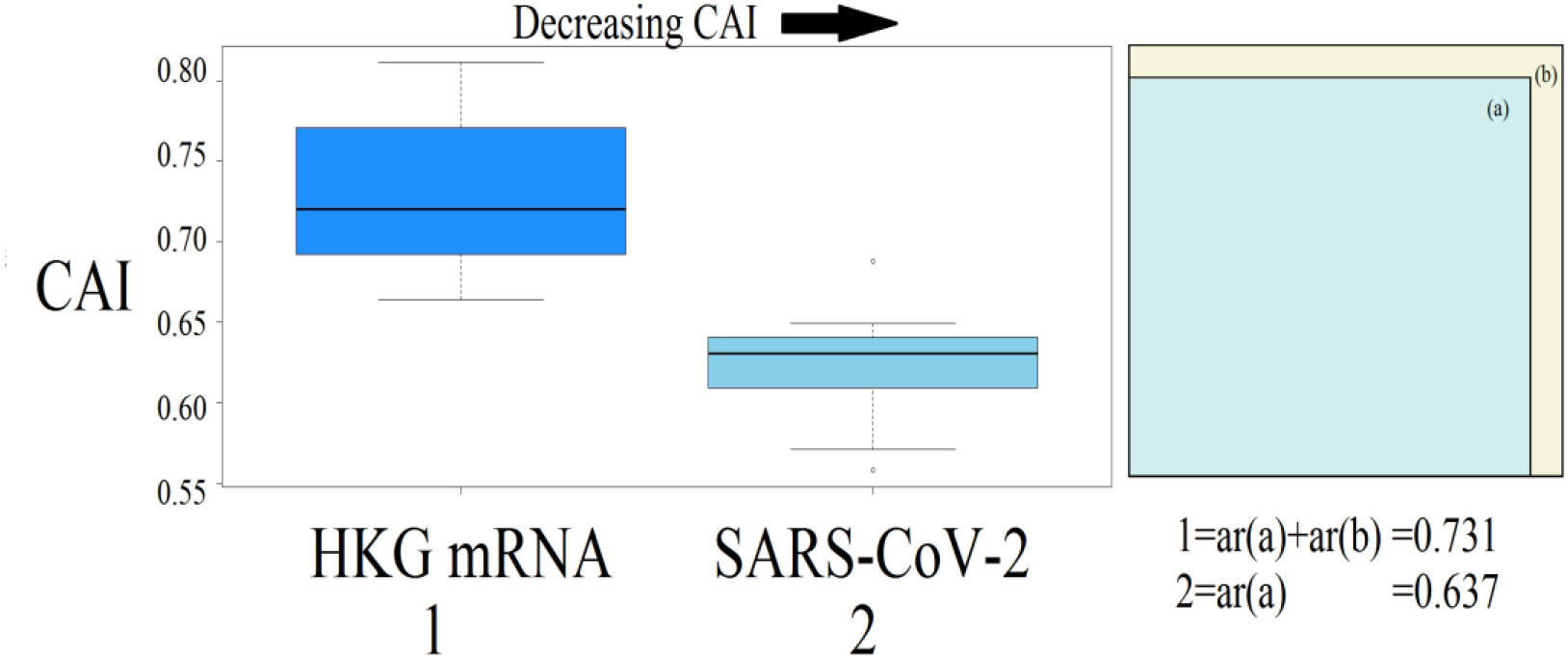
Shows average CAI of HKG and SARS-CoV-2 (predecessor strain) and overlap in the magnitude of their CAI (proxy for gene expression level). Other successor strains have been omitted due to their very close values with the predecessor strain

### 4.5 Investigating the ORF stability of SARS-CoV-2 through mutational simulations

It’s the mRNA stability which is a determiner of the ribosomal abundance during translation. Greater stability of the mRNA results in a higher ribosomal abundance leading to higher translational efficiency(*13*). There is a positive correlation between the ORFs’ stability and CAI in HKG (Refer Results; Table S2.3, Supplementary tables). A similar behavior is seen in SARS-CoV-2 which implies that with increasing stability of the ORFs the gene expression level increases. It was seen that SARS-CoV-2 holds a stronger and significant correlation between ORFs’ stability and CAI, compared to the human HKG, which indicates the virus’s capacity of expression to be greater than that of humans.

Viruses have a very high rate of mutation and in order to examine the scope of plausible codon changes, synonymous codon randomization study was carried on the predecessor strain. This was an in-silico study; simulating synonymous mutations (Refer S1.1, Supplementary materials and methods; Results; Table S2.4, Supplementary tables). As the codon composition in each randomization is identical to the predecessor strain, CAI values will also be identical amongst all the randomized sequences and the predecessor strain. Nonetheless, nucleotide positions have been swapped in the randomized sequences and they will show changes in their folding free energies that will influence the secondary structures and hence the ribosomal abundance. Previous study has established that native mRNAs have the highest stability compared to their in-silico randomized variants (*37*). On the contrary we found that 50% of the synonymously randomized ORFs exhibited folding free energy maxima greater than their native ORFs (Figure 3). We tested for the significance of the difference between the initial (native) and stability maxima for the randomized sequences through t-test and found it to be significant (Refer Results). Thus, the in-silico randomizations of the sequences provide an insight into the mutational possibilities of the virus under the inherent codon frequency. Hence, the virus can develop synonymous polymorphisms without affecting the protein sequences and yet attain ORF stability greater than its predecessor strains. This result in general points the remarkable mutational capability of SARS-CoV-2.

**Figure3:**
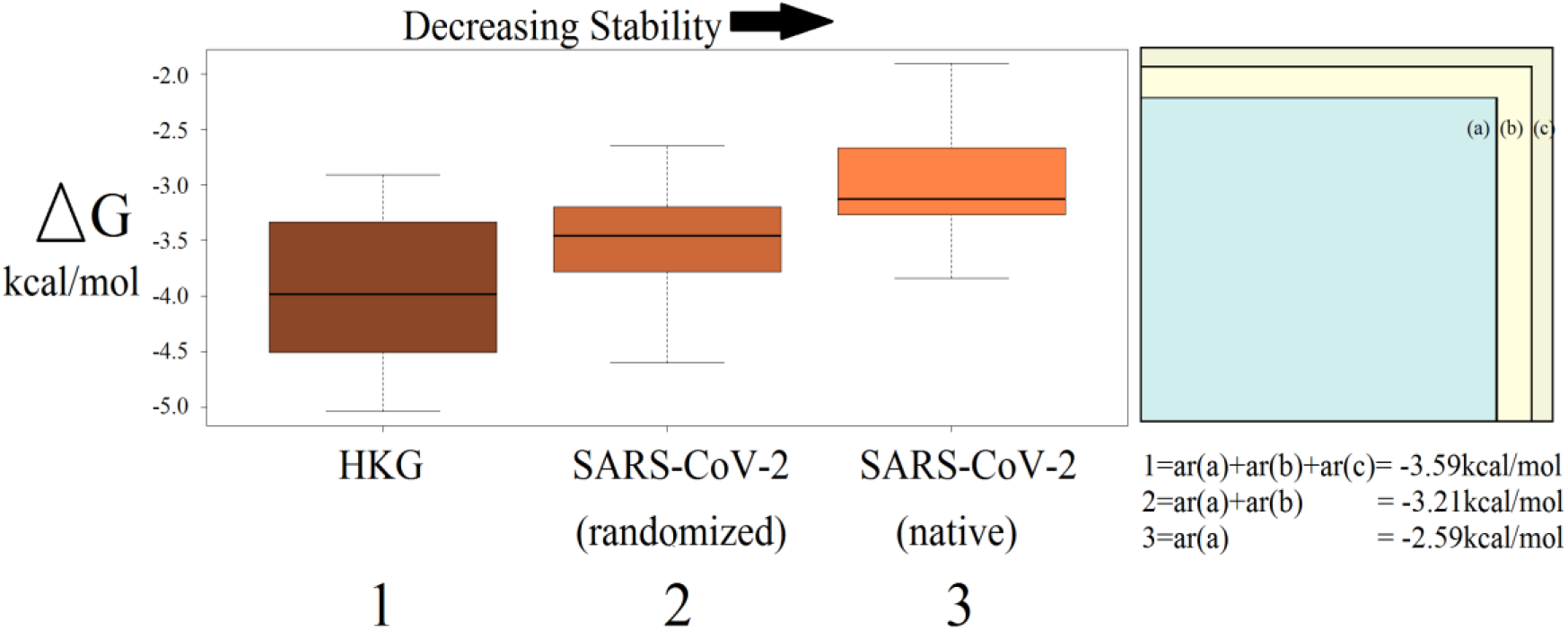
Averages of folding free energies of HKG, SARS-CoV-2 (predecessor strain), SARS-COV-2 (randomized predecessor strain) and overlap in the magnitude of the ΔGs.

### 4.6 Polymorphism and the t-RNA dependence of SARS-CoV-2

As already stated, even under the absence of anti-codons, translation doesn’t stall, instead near-cognate t-RNAs containing anti-codons are employed. In the predecessor strain the t-RNA dependence for the absentee anticodons is seen to be inclined towards a higher gene copy number of near-cognate t-RNAs containing the anticodons. If the viral genome always tends to select anticodons with higher copy number of cognate or near-cognate t-RNAs then, in all the mutations it would show the same. Initially, on screening the successor strains, we found that out of the twelve mutations four were for codons which had no gene for cognate-tRNA. But, after the near cognate t-RNA mapping they all exhibited a choice for the highest gene copy number of near-cognate t-RNA containing the anti-codons. Three out of the twelve mutations were for a higher number of cognate t-RNA, two showed no change and only three preferred, cognate anticodons with a marginally lower t-RNA copy number (Table 2). The result clearly states that, at the genomic level SARS-CoV-2 is thoroughly tuned to be highly compatible with the translational machinery of its host i.e. *Homo sapiens.*

## 5. Conclusion

The findings presented here reveal a strong universal connection between the codon usage patterns in the SARS-CoV-2 with *Homo sapiens* t-RNA pool. Even with a high mutation rate the choice of the codons has a strong inclination for the high copy number of the cognate and near-cognate tRNA containing anticodons. The study highlights that under the conditions of synonymous mutations if the inherent codon composition remains intact, the transcripts have the possibility of gaining greater stability. Generally, with increased stability higher gene expression is seen (*13*). It is interesting to find that the virus even with a higher bias than the human HKG has a considerable codon optimization with the latter. As bias is not the only essential criteria for heightened gene expression the virus has notably balanced both, bias and optimization. We also found that the virus has undergone purifying selection and are exhibiting polymorphism with changes in its ORF1ab polyprotein, surface glycoprotein, and nucleocapsid phosphoprotein. These mutations have been in a very short period of time (~30 days) hence, shows a rapid rate of mutation in the virus. These set of proteins are important for the virus and changes in them might be important for the increased survivability of the virus (*38*). With the simulation study of mutations, rate of mutation in the virus (in ~30 days) and looking at the current global pandemism it’s indicative that SARS-CoV-2 might undergo greater number of mutations producing very high number of variants in geographically different populations of humans.

## Availability of data material

All data were obtained from publicly available databases (mentioned in the ‘Materials and methods’ section) are freely available online. The datasets used and/or analyzed during the current study are also available from the corresponding author on reasonable request.

## Authors’ contributions

M.P.V. designed the study. M.P.V. and T.C.G performed the analysis. MPV, RD drafted the manuscript. M.P.V, R.D. and T.C.G completed the final version of the manuscript. All authors read and approved the manuscript

## Acknowledgement

The authors thank Dr. Debarun Acharya, Dr. Sandip Chakraborty, Dr. Tina Begun and last but not the least Mr. Victor Francis for their immense support

