## Supplementary Information for "An in-silico study on SARS-CoV-2: Its compatibility with human tRNA pool, and the polymorphism arising in a single lineage over a month"

### S1. Supplementary materials and methods:

#### *S1.1. Randomization of the SARS-CoV-2 CDS*

Here, we have used a new randomization process. Synonymous codons occurring in the genes are grouped under respective amino-acid. For every amino-acid in each gene, its position and frequency is noted. During randomization, from the aforementioned group of synonymous codons, a codon is randomly selected for each amino-acid in the native sequence and is used to generate the randomized nucleotide sequences. This selected codon isn't replaced in the group of synonymous codons. It is similar to the process of sampling without replacement. The codon frequency and proteins are identical to native-sequence (Refer Figure S3.1, Supplementary figure). In house python script was written for this process.

#### *S1.2. Minimum folding energy calculation*

Considering the computational cost of processing, ribosomal occupancy of ~30 nucleotides (*1*) and short range interactions of nucleotides for the formation of local secondary structures (*2, 3*) we generated mRNA segments (window) of 50 nucleotides(*2*) and a step size of 30 nucleotides(Temp=37C) (Refer S3.2, Supplementary figure). For each segment, seqfold (*4*) based on various folding algorithms(*5-10*) was used to find the minimum folding energy (ΔG). For each CDS the mean ΔG (kcal/mol) was calculated over the sum total of ΔGs and the number of segments generated. These mean values do not state either exact or approximate estimates for the minimum folding energies of the CDS, instead they are an appropriate indicator of stabilities for comparative analysis. In-house python script was written to generate the mRNA segments.

#### *S1.3. Statistical Calculation*

Choice of statistical tests on small samples tends to resort on non-parametric methods. But, rank methods are least effective on small samples as they fail to give significant outcomes irrespective of the data space(*11*). At this point its intuitive to think that small sample space are more susceptible to sampling errors which might lead to biased conclusions if one has to make statements regarding target population (*12*). Smaller samples obviously have a tendency to generate less precise estimates because the result from an individual study may diverge from the true (population) value. Nevertheless, that doesn't signify an escalated bias. Bias has everything to do with average of the estimates and, only changing the sample size without changing the method for estimations do not affect the average value (*13*). With larger samples the statistic converges more quickly on the true value but this has everything to do with precision (method dependent) and not bias.

The inclination towards non-parametric methods arise due to the inability to assess the distribution shape when there are very few observations, but assuming a sample space lacking normal distribution on the account of scarce data points is an insignificant assumption (*11, 14*). Rather than relying only on analyzing the data size, it's extremely necessary to rely on the similarity of the trends with statistical observations found in already established inferences (*11*). It's essential to have confidence intervals in the former observations for it to be in harmony with the latter (*15*).

### S2. Supplementary tables

*Table S2.1* SARS-CoV-2 and *Homo sapiens* (house-keeping) gene names and accession numbers

| **Accession number** | **Gene Name Human(HKG)** |
|---|---|
| NM_001101.5 | actin beta (ACTB) |
| NM_001289746.2 | glyceraldehyde-3-phosphate dehydrogenase (GAPDH) |
| NM_000291.4 | phosphoglycerate kinase 1 (PGK1) |
| NM_012423.4 | ribosomal protein L13a (RPL13A) |
| NM_053275.3 | ribosomal protein lateral stalk subunit P0 (RPLP0) |
| XM_005254549.3 | beta-2-microglobulin (B2M) |
| NM_004168.4 | succinate dehydrogenase complex flavoprotein subunit A (SDHA) |
| NM_000181.4 | glucuronidase beta (GUSB) |
| NM_000190.4 | hydroxymethylbilane synthase (HMBS) |
| NM_000194.3 | hypoxanthine phosphoribosyltransferase 1 (HPRT1) |
| NM_003194.5 | TATA-box binding protein (TBP) |
| NM_004542 | NADH:ubiquinone oxidoreductase subunit A3 (NDUFA3) |
| **Accession number (SARS-CoV-2)** | **Date** |
| MT539164 (predecessor) | 2020-05-30 |
| MT675933 (successor) | 2020-06-29 |
| MT675937 (successor) | 2020-06-29 |
| MT675938 (successor) | 2020-06-29 |
| MT675939 (successor) | 2020-06-29 |
| MT675940 (successor) | 2020-06-29 |
| MT675941 (successor) | 2020-06-29 |

*Table S2.2 Homo sapiens* near-cognate t-RNA mapping with SARS-CoV-2 codons

| **Amino-Acid** | **Codon_SARS-CoV-2** | **Anti-codon** | **t-RNA count by Anti-codons(*Homo sapiens*)** | **t-RNA mapping** | **t-RNA count** |
|---|---|---|---|---|---|
| Ala | GCA | TGC | 8 | TGC | 8 |
| Ala | GCC | GGC | 0 | **AGC** | **22** |
| Ala | GCG | CGC | 4 | CGC | 4 |
| Ala | GCT | AGC | 22 | AGC | 22 |
| Cys | TGC | GCA | 29 | GCA | 29 |
| Cys | TGT | ACA | 0 | **GCA** | **29** |
| Asp | GAC | GTC | 13 | GTC | 13 |
| Asp | GAT | ATC | 0 | **GTC** | **13** |
| Glu | GAA | TTC | 7 | TTC | 7 |

| Amino-Acid | Codon_SARS-CoV-2 | Anti-codon | t-RNA count by Anti-codons(*Homo sapiens*) | t-RNA mapping | t-RNA count |
|---|---|---|---|---|---|
| Glu | GAG | CTC | 8 | CTC | 8 |
| Phe | TTC | GAA | 10 | GAA | 10 |
| Phe | TTT | AAA | 0 | **GAA** | **10** |
| Gly | GGA | TCC | 9 | TCC | 9 |
| Gly | GGC | GCC | 14 | GCC | 14 |
| Gly | GGG | CCC | 5 | CCC | 5 |
| Gly | GGT | ACC | 0 | **GCC** | **14** |
| His | CAC | GTG | 10 | GTG | 10 |
| His | CAT | ATG | 0 | **GTG** | **10** |
| Ile | ATA | TAT | 5 | TAT | 5 |
| Ile | ATC | GAT | 3 | GAT | 3 |
| Ile | ATT | AAT | 14 | AAT | 14 |
| Lys | AAA | TTT | 12 | TTT | 12 |
| Lys | AAG | CTT | 15 | CTT | 15 |
| Leu | CTA | TAG | 3 | TAG | 3 |
| Leu | CTC | GAG | 0 | **AAG** | **9** |
| Leu | CTG | CAG | 9 | CAG | 9 |
| Leu | CTT | AAG | 9 | AAG | 9 |
| Leu | TTA | TAA | 4 | TAA | 4 |
| Leu | TTG | CAA | 6 | CAA | 6 |
| Met | ATG | CAT | 10 | CAT | 10 |
| Asn | AAC | GTT | 20 | GTT | 20 |
| Asn | AAT | ATT | 0 | **GTT** | **20** |
| Pro | CCA | TGG | 7 | TGG | 7 |
| Pro | CCC | GGG | 0 | **AGG** | **9** |
| Pro | CCG | CGG | 4 | CGG | 4 |
| Pro | CCT | AGG | 9 | AGG | 9 |
| Gln | CAA | TTG | 6 | TTG | 6 |
| Gln | CAG | CTG | 13 | CTG | 13 |
| Arg | AGA | TCT | 6 | TCT | 6 |
| Arg | AGG | CCT | 5 | CCT | 5 |
| Arg | CGA | TCG | 6 | TCG | 6 |
| Arg | CGC | GCG | 0 | **ACG** | **7** |
| Arg | CGG | CCG | 4 | CCG | 4 |
| Arg | CGT | ACG | 7 | ACG | 7 |
| Ser | AGC | GCT | 8 | GCT | 8 |
| Ser | AGT | ACT | 0 | **GCT** | **8** |
| Ser | TCA | TGA | 4 | TGA | 4 |
| Ser | TCC | GGA | 0 | **AGA** | **9** |
| Ser | TCG | CGA | 4 | CGA | 4 |
| Ser | TCT | AGA | 9 | AGA | 9 |

| Amino-Acid | Codon_SARS-CoV-2 | Anti-codon | t-RNA count by Anti-codons(*Homo sapiens*) | t-RNA mapping | t-RNA count |
|---|---|---|---|---|---|
| Thr | ACA | TGT | 6 | TGT | 6 |
| Thr | ACC | GGT | 0 | **AGT** | **9** |
| Thr | ACG | CGT | 5 | CGT | 5 |
| Thr | ACT | AGT | 9 | AGT | 9 |
| Val | GTA | TAC | 5 | TAC | 5 |
| Val | GTC | GAC | 0 | **AAC** | **9** |
| Val | GTG | CAC | 11 | CAC | 11 |
| Val | GTT | AAC | 9 | AAC | 9 |
| Trp | TGG | CCA | 7 | CCA | 7 |
| Tyr | TAC | GTA | 13 | GTA | 13 |
| Tyr | TAT | ATA | 0 | **GTA** | **13** |

*Zero denotes the absence of cognate t-RNA. Codons and anti-codons are in 5’ → 3’

*Table S2.3* CAI and folding free energy values for SARS-CoV-2and HKG

| Accession number | Gene_Name SARS-CoV-2 | CAI | ΔG(kcal/mol) |
|---|---|---|---|
| MT539164 | envelope protein | 0.558 | -1.91 |
| | ORF10 protein | 0.571 | -3.17 |
| | ORF6 protein | 0.606 | -2.62 |
| | ORF7b | 0.612 | -2 |
| | ORF8 protein | 0.628 | -2.73 |
| | membrane glycoprotein | 0.629 | -3.3 |
| | ORF3a protein | 0.632 | -3.63 |
| | ORF1a polyprotein | 0.634 | -3.24 |
| | ORF1b polyprotein | 0.635 | -3.16 |
| | surface glycoprotein | 0.646 | -3.1 |
| | ORF7a protein | 0.649 | -2.72 |
| | nucleocapsid phosphoprotein | 0.688 | -3.84 |
| **Accession number** | **Gene Name Human(HKG)** | **CAI human** | **ΔG(kcal/mol)** |
| NM_001101.5 | actin beta (ACTB) | 0.811 | -3.82 |
| NM_001289746.2 | glyceraldehyde-3-phosphate dehydrogenase (GAPDH) | 0.7 | -4.8 |
| NM_000291.4 | phosphoglycerate kinase 1 (PGK1) | 0.7 | -3.01 |
| NM_012423.4 | ribosomal protein L13a (RPL13A) | 0.666 | -3.66 |
| NM_053275.3 | ribosomal protein lateral stalk subunit P0 (RPLP0) | 0.771 | -5.04 |
| XM_005254549.3 | beta-2-microglobulin (B2M) | 0.664 | -2.91 |
| NM_004168.4 | succinate dehydrogenase complex flavoprotein subunit A (SDHA) | 0.745 | -3.88 |
| NM_000181.4 | glucuronidase beta (GUSB) | 0.785 | -4.22 |
| NM_000190.4 | hydroxymethylbilane synthase (HMBS) | 0.77 | -4.88 |
| NM_000194.3 | hypoxanthine phosphoribosyltransferase 1 (HPRT1) | 0.684 | -2.93 |
| NM_003194.5 | TATA-box binding protein (TBP) | 0.72 | -4.09 |
| NM_004542 | NADH:ubiquinone oxidoreductase subunit A3 (NDUFA3) | 0.758 | -4.2 |

*Table S2.4* The folding free energies of HKG, SARS-CoV-2 and ΔG maxima amongst its randomized sequences

| **MT539164** | **Native ΔG** | **Randomized ΔG Maxima** | **HKG ΔG** |
|---|---|---|---|
| nucleocapsid phosphoprotein | -3.84 | -3.55 | -5.04 |
| ORF3a protein | -3.63 | -2.9 | -4.88 |
| membrane glycoprotein | -3.3 | -3.22 | -4.8 |
| ORF1a polyprotein | -3.24 | -2.44 | -4.22 |
| ORF10 protein | -3.17 | -4.6 | -4.2 |
| ORF1b polyprotein | -3.16 | -2.39 | -4.09 |
| surface glycoprotein | -3.1 | -2.79 | -3.88 |
| ORF8 protein | -2.73 | -3.94 | -3.82 |
| ORF7a protein | -2.72 | -3.73 | -3.66 |
| ORF6 protein | -2.62 | -3.48 | -3.01 |
| ORF7b | -2 | -2.65 | -2.93 |
| envelope protein | -1.91 | -3.43 | -2.91 |

*Table S2.5* Mutations and their positions in the different ORFs of SARS-CoV-2 with respect to MT539164 (predecessor strain) (AA=Amino-Acid)

| Annotation | Name | Initial Codons | Mutated Codons | Position | Initia AA | Mutated AA |
|---|---|---|---|---|---|---|
| **Accession Number** | | | | | | |
| MT675933 | ORF1a polyprotein | CAG | CCG | 2026 | gln | pro |
| | | TGC | TGT | 2569 | cys | cys |
| | | GTG | GTT | 4033 | val | val |
| | | TTT | TTC | 11353 | phe | phe |
| | ORF1b polyprotein | CAG | CCG | 2026 | gln | pro |
| | | TGC | TGT | 2569 | cys | cys |
| | | GTG | GTT | 4033 | val | val |
| | surface glycoprotein | TTG | TTT | 160 | leu | phe |
| | | GAC | GAT | 880 | asp | asp |
| | nucleocapsid phosphoprotein | TCA | TTA | 580 | ser | leu |
| **Accession Number** | | | | | | |
| MT675937 | ORF1a polyprotein | CAG | CCG | 2026 | gln | pro |
| | | TGC | TGT | 2569 | cys | cys |
| | | GTG | GTT | 4033 | val | val |

| Annotation | Name | Initial Codons | Mutated Codons | Position | Initia AA | Mutated AA |
|---|---|---|---|---|---|---|
| | | TTT | TTC | 11353 | phe | Phe |
| | ORF1b polyprotein | TTA | TTG | 16246 | leu | leu |
| | | CTC | TTC | 18304 | leu | phe |
| | surface glycoprotein | TTG | TTT | 160 | leu | phe |
| | | GAC | GAT | 880 | asp | asp |
| | nucleocapsid phosphoprotein | TCA | TTA | 580 | ser | leu |
| **Accession Number** | | | | | | |
| MT675938 | ORF1a polyprotein | CAG | CCG | 2026 | gln | pro |
| | | TGC | TGT | 2569 | cys | cys |
| | | GTG | GTT | 4033 | val | val |
| | | TTT | TTC | 11353 | phe | phe |
| | ORF1b polyprotein | TTA | TTG | 16246 | leu | leu |
| | | CTC | TTC | 18304 | leu | phe |
| | | ACA | ATA | 18889 | thr | ile |
| | surface glycoprotein | TTG | TTT | 160 | leu | phe |
| | | GAC | GAT | 880 | asp | asp |
| | nucleocapsid phosphoprotein | TCA | TTA | 580 | ser | leu |
| **Accession Number** | | | | | | |
| MT675939 | ORF1a polyprotein | CAG | CCG | 2026 | gln | pro |
| | | TGC | TGT | 2569 | cys | cys |
| | | GTG | GTT | 4033 | val | val |
| | | TTT | TTC | 11353 | phe | phe |
| | ORF1b polyprotein | CTT | CCT | 14143 | leu | pro |
| | | CTT | TTT | 15088 | leu | phe |
| | | TTA | TTG | 16246 | leu | leu |
| | | CTC | TTC | 18304 | leu | phe |
| | surface glycoprotein | TTG | TTT | 160 | leu | phe |
| | | GAC | GAT | 880 | asp | asp |
| | nucleocapsid phosphoprotein | TCA | TTA | 580 | ser | leu |
| **Accession Number** | | | | | | |
| MT675940 | ORF1a polyprotein | CAG | CCG | 2026 | gln | pro |
| | | TGC | TGT | 2569 | cys | cys |

| Annotation | Name | Initial Codons | Mutated Codons | Position | Initia AA | Mutated AA |
|---|---|---|---|---|---|---|
| | | GTG | GTT | 4033 | val | Val |
| | | TTT | TTC | 11353 | phe | phe |
| | ORF1b polyprotein | CTT | CCT | 14143 | leu | pro |
| | | TTA | TTG | 16246 | leu | leu |
| | | CTC | TTC | 18304 | leu | phe |
| | | ACA | ATA | 18889 | thr | ile |
| | surface glycoprotein | TTG | TTT | 160 | leu | phe |
| | | GAC | GAT | 880 | asp | asp |
| | nucleocapsid | TCA | TTA | 580 | ser | leu |
| **Accession Number** | | | | | | |
| MT675941 | ORF1a polyprotein | CAG | CCG | 2026 | gln | pro |
| | | TGC | TGT | 2569 | cys | cys |
| | | GTG | GTT | 4033 | val | val |
| | | TTT | TTC | 11353 | phe | phe |
| | ORF1b polyprotein | CTT | CCT | 14143 | leu | pro |
| | | CTT | TTT | 15088 | leu | phe |
| | | TTA | TTG | 16246 | leu | leu |
| | | CTC | TTC | 18304 | leu | phe |
| | surface gylcoprotein | TTG | TTT | 160 | leu | phe |
| | | GAC | GAT | 880 | asp | asp |
| | nucleocapsid phosphoprotein | TCA | TTA | 580 | ser | leu |

### S3. Supplementary figures

*Figure S3.1* Schematic representation of creating mRNAs through randomization of the synonymous codons within each mRNA. The inherent codon frequency and amino-acid sequences are identical to the native sequences. The numbers of swappings in each randomization are higher than depicted in the figure

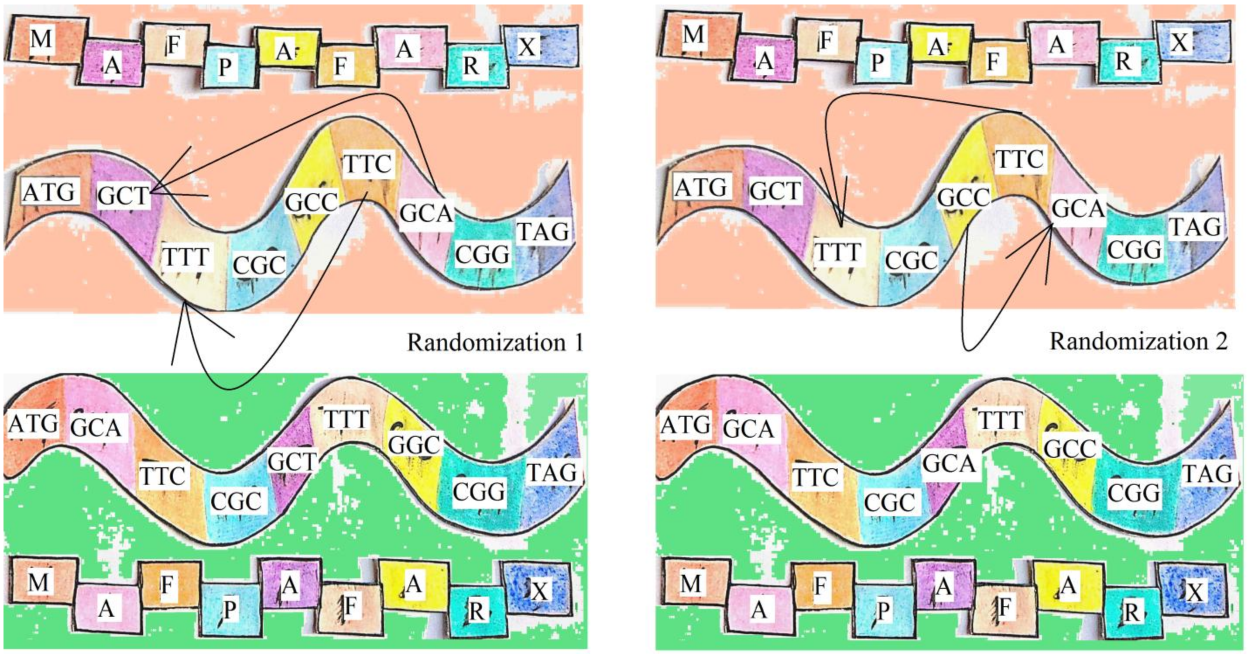


*Figure S3.2* Schematic representation for the selection of window size for the calculation of mRNA stability based on the ribosomal occupancy (~ 30 nucleotides) of the nucleotides during translation

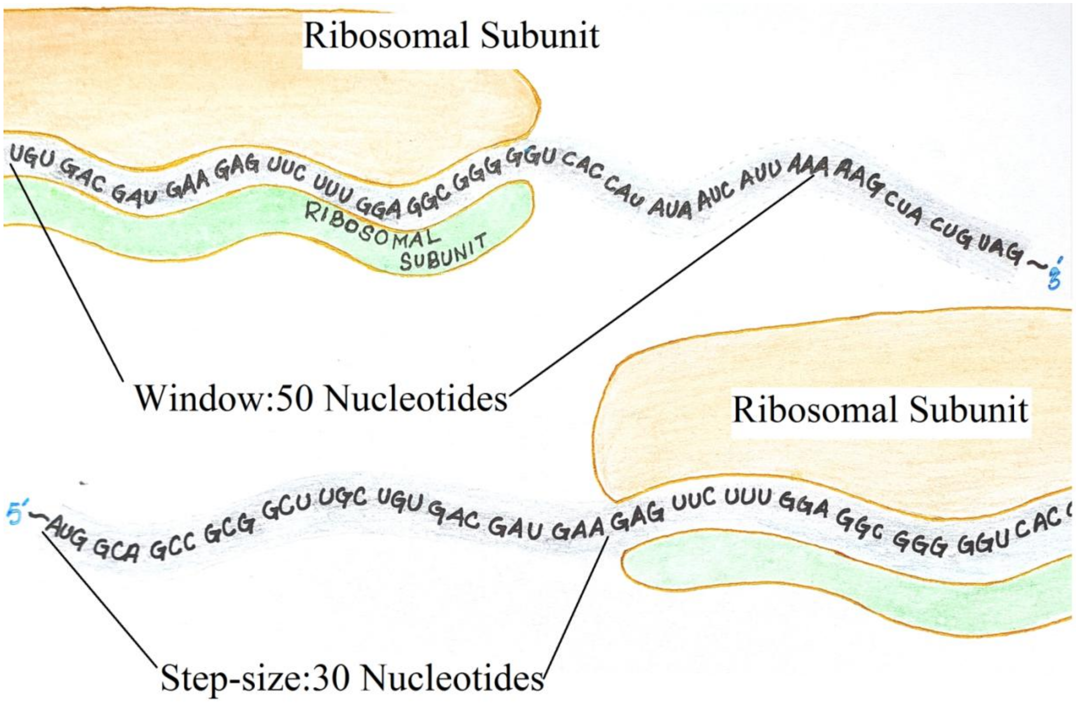

Supplementary Information References:

1. N. T. Ingolia, S. Ghaemmaghami, J. R. Newman, J. S. Weissman, Genome-wide analysis in vivo of translation with nucleotide resolution using ribosome profiling. *Science* **324**, 218-223 (2009).
2. C. Park, X. Chen, J. R. Yang, J. Zhang, Differential requirements for mRNA folding partially explain why highly expressed proteins evolve slowly. *Proc Natl Acad Sci U S A* **110**, E678-686 (2013).
3. M. J. Doktycz, F. W. Larimer, M. Pastrnak, A. Stevens, Comparative analyses of the secondary structures of synthetic and intracellular yeast MFA2 mRNAs. *Proc Natl Acad Sci U S A* **95**, 14614-14621 (1998).
4. J. Timmons , l. Leshane. (M.I.T, 2019), pp. Predict the minimum free energy structure of nucleic acids.
5. R. Nussinov, A. B. Jacobson, Fast algorithm for predicting the secondary structure of single-stranded RNA. *Proc Natl Acad Sci U S A* **77**, 6309-6313 (1980).
6. M. Zuker, P. Stiegler, Optimal computer folding of large RNA sequences using thermodynamics and auxiliary information. *Nucleic Acids Res* **9**, 133-148 (1981).
7. J. A. Jaeger, D. H. Turner, M. Zuker, Improved predictions of secondary structures for RNA. *Proc Natl Acad Sci U S A* **86**, 7706-7710 (1989).
8. J. SantaLucia, D. Hicks, The thermodynamics of DNA structural motifs. *Annu Rev Biophys Biomol Struct* **33**, 415-440 (2004).
9. D. H. Turner, D. H. Mathews, NNDB: the nearest neighbor parameter database for predicting stability of nucleic acid secondary structure. *Nucleic Acids Res* **38**, D280-282 (2010).
10. M. Ward, A. Datta, M. Wise, D. H. Mathews, Advanced multi-loop algorithms for RNA secondary structure prediction reveal that the simplest model is best. *Nucleic Acids Res* **45**, 8541-8550 (2017).
11. J. M. Bland, D. G. Altman, Analysis of continuous data from small samples. *BMJ* **338**, a3166 (2009).
12. A. Hackshaw, Small studies: strengths and limitations. *Eur Respir J* **32**, 1141-1143 (2008).
13. L. Lin, Bias caused by sampling error in meta-analysis with small sample sizes. *PLoS One* **13**, e0204056 (2018).

14. C. J. Baker, D. L. Kasper, M. S. Edwards, G. Schiffman, Influence of preimmunization antibody levels on the specificity of the immune response to related polysaccharide antigens. *N Engl J Med* **303**, 173-178 (1980).
15. B. Weaver, R. Koopman, An SPSS Macro to Compute Confidence Intervals for Pearson's Correlation. *The Quantitative Methods for Psychology* **10**, 29-39 (2014).
